# How human behavior drives the propagation of an emerging infection: the case of the 2014 Chikungunya outbreak in Martinique

**DOI:** 10.1101/052183

**Authors:** Benjamin Roche, Béatrice Gaillard, Lucas Léger, Renélise Moutenda, Thomas Sochacki, Bernard Cazelles, Martine Ledrans, Alain Blateau, Didier Fontenille, Manuel Etienne, Frédéric Simard, Marcel Salathé, André Yébakima

## Abstract

Understanding the spatio-temporal dynamics of endemic infections is of critical importance for a deeper understanding of pathogen transmission, and for the design of more efficient public health strategies. However, very few studies in this domain have focused on emerging infections, generating a gap of knowledge that hampers epidemiological response planning. Here, we analyze the case of a Chikungunya outbreak that occurred in Martinique in 2014. Using time series estimates from a network of sentinel practitioners covering the entire island, we first analyze the spatio-temporal dynamics and show that the largest city has served as the epicenter of this epidemic. We further show that the epidemic spread from there through two different propagation waves moving northwards and southwards, probably by individuals moving along the road network. We then develop a mathematical model to explore the drivers of the temporal dynamics of this mosquito-borne virus. Finally, we show that human behavior, inferred by a textual analysis of messages published on the social network Twitter, is required to explain the epidemiological dynamics over time. Overall, our results suggest that human behavior has been a key component of the outbreak propagation, and we argue that such results can lead to more efficient public health strategies specifically targeting the propagation process.

## Introduction

It is well known that for most infectious diseases, transmission intensity fluctuates through space and time (1). Understanding the spatio-temporal dynamics of infectious diseases provides considerable insights into our understanding of their epidemiology (1–4) and can help design better strategies for their control (5, 6). The transmission dynamics of numerous endemic pathogens, ranging from childhood diseases (3) to vector-borne diseases (7), have been studied in many different countries (8, 9). As a result, numerous factors have been identified that drive the temporal dynamics of pathogen transmission (10), such as the abundance of vector population for vector-borne diseases (11), or abiotic factors, which can drive the dynamics of environmentally-transmitted agents like cholera (12). They also include host behavior, such as the switch between school and holiday periods that shapes seasonality of childhood diseases (13), or more complex social mechanisms, involving belief-based or prevalence-based feelings (14).

Local transmission can also be influenced by movements of infectious individuals between cities, especially in small localities and islands (15). This leads to a spatial structure of pathogen spread which has been extensively documented for many communicable diseases (1, 7). A pattern of travelling waves from large cities to rural areas has been highlighted for childhood diseases in the UK (3) or Senegal (16), as well as for vector-borne diseases such as Dengue fever in Thailand (7, 17).

Although these temporal and spatio-temporal patterns have been documented for many endemic diseases, studies on emerging infections remain rare, creating a large information gap (but see (18) for a recent example). This gap is detrimental to public health because an understanding of the initial stages of propagation is highly relevant for the overall disease epidemiology. Observations during the early stages of an epidemic allow public health authorities to adapt their strategies quickly (19) and to efficiently organize a control response in the face of such highly unpredictable events. An efficient early response is particularly crucial in the case of emerging infections where most of the population is assumed to be susceptible due to the absence of pre-existing immunity.

The recurrence of these emerging events are currently threatening much of the progress made by public health campaigns during the past decades (20, 21), making a solid understanding of early-stage epidemic dynamics a clear research priority. To this extent, the temporal and spatio-temporal dynamics of Chikungunya outbreak in Martinique Island represents a unique semi-natural experimental case study. Introduced in December 2013 in Caribbean Islands (22) and in Martinique (23), the Chikungunya virus, transmitted by the mosquito *Aedes aegypti* in this area, has spread throughout the whole island, resulting in more than 72,500 infections and 51 deaths (24). Since the island has a relatively small land area (1,040 km^2^), introductions mainly happen through the largest cities where the harbor and airport are located, allowing us to study the natural drivers of the spatio-temporal propagation without the scrambling effect of multiple immigration routes.

We aimed to identify the main drivers of the temporal and spatio-temporal dynamics of the 2014 Chikungunya outbreak in Martinique. We first analyzed the similarity between epidemiological time series recorded within each locality to quantify the propagation through space and time. Focusing on the period preceding a large spatial propagation throughout the island, we then fitted a mathematical model to quantify the contribution of three potential drivers, namely (i) mosquito abundance (inferred from entomological surveys), (ii) awareness of the epidemic in the local human population, and (iii) the interest for protection against the disease (the latter two both inferred from a textual analysis of messages recorded on the social network Twitter) on the temporal dynamics of the outbreak. Based on the identified importance of human behavior during this outbreak, we argue that more research should focus on quantitative assessment of human behavior in the context of emerging infections, and that this component should be considered very seriously in epidemiological response planning in the face of unexpected disease outbreaks.

## Materials and methods

### Epidemiological Data

Our epidemiological data includes the number of new suspected Chikungunya cases within each locality for each week during the first nine months of the outbreak (from December 2013 to August 2014). Here, we use the number of suspected cases rather than the number of confirmed cases because biological confirmation has not been made routinely after the first 5 months owing to logistic complexity, and thus does not reflect the disease activity over the whole period. These incidence time series have been estimated based on a network of sentinel practitioners that covers all the localities on the island with at least one medical doctor, resulting in 28 localities considered (over a total number of 34 localities throughout the island). Each week, the total number of suspected cases collected by sentinel practitioners is extrapolated to the whole island using the ratio “medical activity of sentinel practitioners present during the week” / “medical activity of all the practitioners in Martinique” (25). These time series have been smoothened through Fast-Fourrier Transform (FFT algorithm (26)) in order to remove excessive stochasticity.

### Entomological data

Mosquito abundance is expected to play a significant role in pathogen transmission rate. Taking advantage of long term surveys that have been conducted in Martinique during the past 15 years, we modeled statistically the abundance of mosquitoes through time, assuming that this abundance is linked to the proportion of infested houses visited during routine surveillance, and to the probability of mosquito presence. More precisely, we quantify the contribution of the different climatic and land-use data available through Generalized Linear Mixed models (27) in order to explain the probability of mosquito presence. This step allows us to derive a robust estimate of the population dynamics of *Aedes aegypti* through time (see details in Supplementary Materials S1).

### Human behavior data

Because insecticides are expected to have a limited efficiency according to the high level of mosquito resistance observed in Martinique (28), we aimed to infer the efficiency of intensive communication campaigns aimed at the public, operated by local public health authorities. To do that, we assume that human behavior that could impact pathogen transmission – i.e. through larval source reduction (e.g. removal of stagnant water) and/or individual protection (use of repellents, bednets, etc.) against mosquitoes – can be estimated by the amount of relevant messages posted by local residents on the online social network Twitter talking about the outbreak. We quantified the awareness of the epidemic through the number of messages posted on Twitter during the first 9 months of the outbreak that contained the word *Chikungunya*. In order to account solely for sentiments from people who were located in Martinique during the outbreak, we considered only messages by Twitter users who declared to be based in Martinique. Then, we quantified the protection behavior through the presence of a sentiment of protection need expressed in the tweets. We textually analyzed the content of each of the 423 tweets messages recorded during this period (including re-tweets) to identify 73 tweets messages with the presence of this feeling (still including re-tweets). Messages classification is detailed in Supplementary Materials (section S2).

### Spatio-temporal analysis

The spatio-temporal dynamics were first visually assessed by plotting the scaled incidence rate for each locality through time. This visual exploration allowed formulating a hypothesis about the structure of the spatio-temporal dynamics that were then tested statistically. Following previous works on the metapopulation of infectious diseases (17, 29), we tested if the similarity between times series of incidence rate at each locality was associated with the geographic distance between these localities. If a travelling pattern exists, the similarity between times series is expected to decrease with geographic distance between them (29). To quantify the similarity between time series, we used (i) the Euclidean distance at each time step of the time series over the whole time series and (ii) at the epidemic peak (the week when the locality has reached its maximal incidence rate).

### Temporal analysis

To quantify the contribution of each of the potential drivers of temporal dynamics, we used a mathematical model framed within the SIR framework (30):

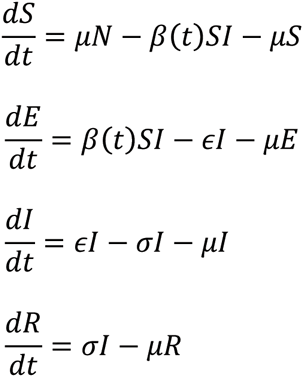

where *S* represents susceptible individuals that are not infected. These individuals can become exposed (*E*) at a rate *β*(*t*). After a latency period of 1/ε (asssumed here to be 3 days(31)), exposed individuals become infectious (*I*) and then infect susceptible mosquitoes, which can in turn infect susceptible humans. Finally, infectious individuals recover at rate *σ* (assumed to be 4 days^−1^ ˙ind^−1^, (31)) and are then permanently immunized against the disease (*R*). Birth and death rates are identical (*μ*) in order to keep population size constant. Since we focus on an emerging event over a short period of time, we assume that human demography does not play a significant role in the epidemic dynamics. We include the contribution of the three potential temporal drivers through a fluctuating forced transmission rate:

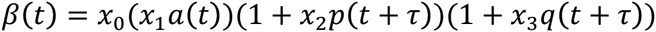

where *x*_*0*_ is the average transmission rate through time, (*x*_1_*a*(*t*)) represents the impact of mosquito population dynamics on transmission rate (*x*_*1*_ is a constant and *a*(*t*) is the estimated mosquito abundance at time t), (1 + *x*_2_*p*(*t*)) represents the influence of protection applied by human population, as measured by the textual analysis of tweets (*x*_*2*_ is a constant and *p(t)* represents the normalized activity on social network expressing a need for protection) and (1 + *x*_3_*q*(*t*)) represents the influence of activity - and therefore of the epidemic awareness - as measured on Twitter ( *x*_*3*_ is a constant and *q(t)* represents the normalized overall activity on Twitter discussing the Chikungunya outbreak). The parameter *τ*represents the lag between the impact quantified on Twitter and its consequences for transmission. We have assumed that the behavior measured on Twitter could be a real-time indicator (*τ* = 0), a delayed indicator (*τ* = 1) when individuals talk on Twitter after having applied the protection, or an anticipated indicator (*τ* = −1) when the Twitter activity reflects the behavior change one month before its impact on mosquito population. The *β*(*t*) function is constrained positive. We then looked for the best estimation of *x*_*1*_, *x*_*2*_ and *x*_*3*_ that allows the mathematical model to reproduce the incidence dynamics observed. Technically, we maximized the likelihood of the model predictions knowing the data by assuming a Gaussian distribution of the model errors through the Nelder-Manson algorithm implemented within the *optim* package in R (32). All details are given in supplementary materials S3.

## Results

The first cases have been declared in Fort-De-France, the largest city on the Island (33). The visual analysis of the spatio-temporal dynamics shows two travelling waves from this city, northwards and southwards (Figure 1). Interestingly, in the north of the island, the association between local epidemiological dynamics and geographic distance relies on the timing of epidemic peak (r=0.6454, p-value=0.0094) rather than on the whole series (r=0.49, p-value=0.061), while in the south of the island, it relies on the whole times series (r=0.72, p-value=0.0037) rather than on the epidemic peak (r=0.39, p-value=0.16). This difference will be discussed later. Based on these correlations, we use the correlation coefficients to characterize the invasion sequence of the virus throughout the island (Figure 2).

**Figure 1:**
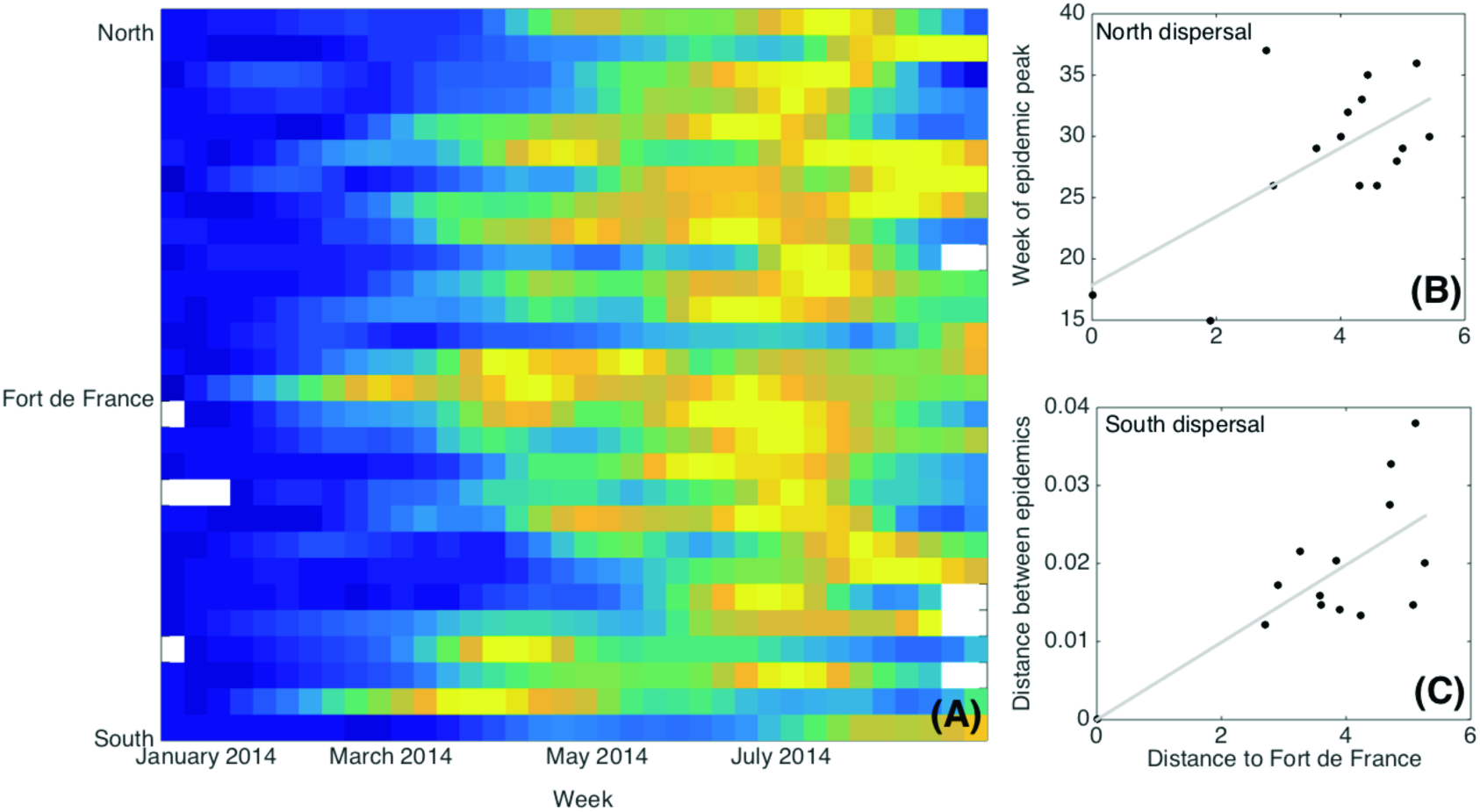
**Summary of the spatio-temporal dynamics of the 2014 Chikungunya outbreak in Martinique. (A) Colors represent the scaled incidence rate (ranking from 0 in dark to 1 in hot yellow) for each locality (rows), ranked from the extreme south to the extreme north of the island according to its position from Fort-De-France, and each week (column). Two waves appear from Fort-De-France, northwards and southwards. Geographic distance (log of km) between different localities through road (x-axis) and (B) the week of the epidemic peak (r=0.6454, p-value=0.0094) and (C) the Euclidean distance between the whole time series (r=0.72, p-value=0.0037).**

**Figure 2:**
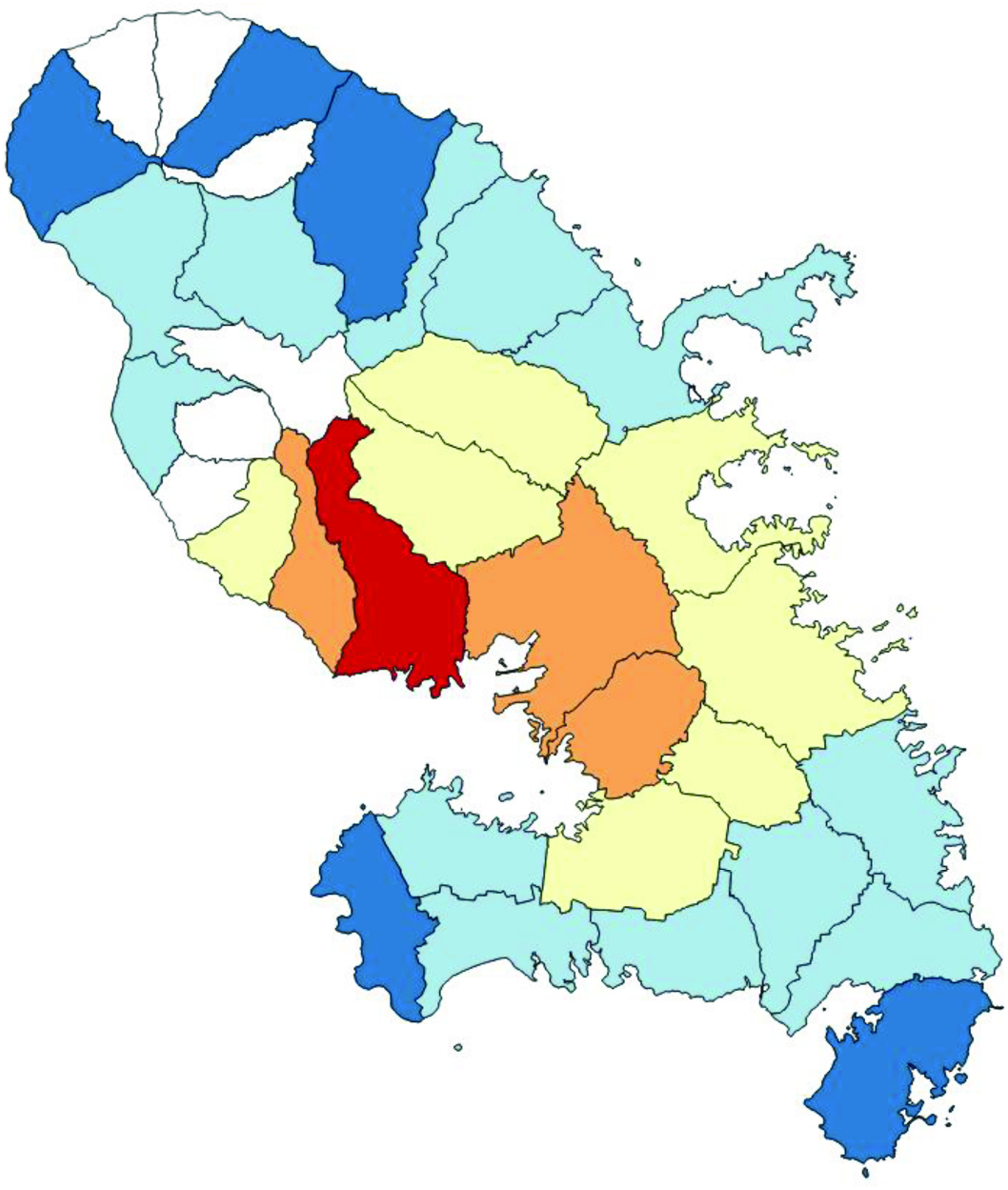
**Invasion sequence of the Chikungunya outbreak based on the correlation between geographic distance and epidemiological dynamics (see main text for more details). Colors follow a gradient from red (epicenter of the epidemics, Fort-de-France) to dark blue (last localities to have been affected, i.e., showing the lowest correlation coefficient).**

In order to focus on a time period when pathogen dispersal was limited, we limit the analysis of the temporal dynamics of the epidemic to the time period before it began spreading to remote areas within the island, *i.e.* from December 2013 to May 2014. We estimate parameters *x*_*1*_, *x*_*2*_ and *x*_*3*_ for all possible combinations of transmission drivers (mosquito abundance, awareness of the epidemic, and the need for protection) and an anticipated (*τ* = −1), real-time (*τ* = 0) or delayed (*τ* = 1) indicator of human behavior impact on transmission. Comparisons between the Mean-Squared Error (using Akaike Information Criterion gives identical results) of each of these models show that the model including both mosquito abundance and the expressed need for protection, with an anticipated notification on Twitter, is the most parsimonious explanation of the temporal dynamics observed (Table 1). Moreover, the model predicts around 28,000 new cases over the six-month period, an estimate in line with the slightly more than 30,000 cases recorded by epidemiological data, highlighting the accuracy of this mathematical model despite its simplicity (Figure 3).

**Table 1:** **Results of model estimation. We show here the squared root of the Mean-Squared Error instead of AIC in order to show the difference between the observed and predicted number of cases. The best model includes the variation in mosquito abundance and the expressed need for protection represented by the proportion of tweets talking about protection against the mosquito in the set of all tweets that included the word *Chikungunya* (only Twitter accounts declared in Martinique have been considered).**

| Parameters included in transmission rate | Twitter as an anticipated indicator ($\tau=-1$) | Twitter as a real-time indicator ($\tau=0$) | Twitter as a delayed indicator ($\tau=1$) |
| --- | --- | --- | --- |
| None | 9088 | 9088 | 9088 |
| Mosquito abundance (MA) | 6980 | 6980 | 6980 |
| Expressed protection need (EPN) | 3897 | 7546 | 5191 |
| Epidemics awareness (EA) | 7797 | 8240 | 6878 |
| MA and EPN | 2402 | 4058 | 3161 |
| MA and EA | 7242 | 8518 | 5484 |
| EPN and EA | 7218 | 7529 | 3685 |
| MA, EPN and EA | 4389 | 7639 | 2675 |

**Figure 3:**
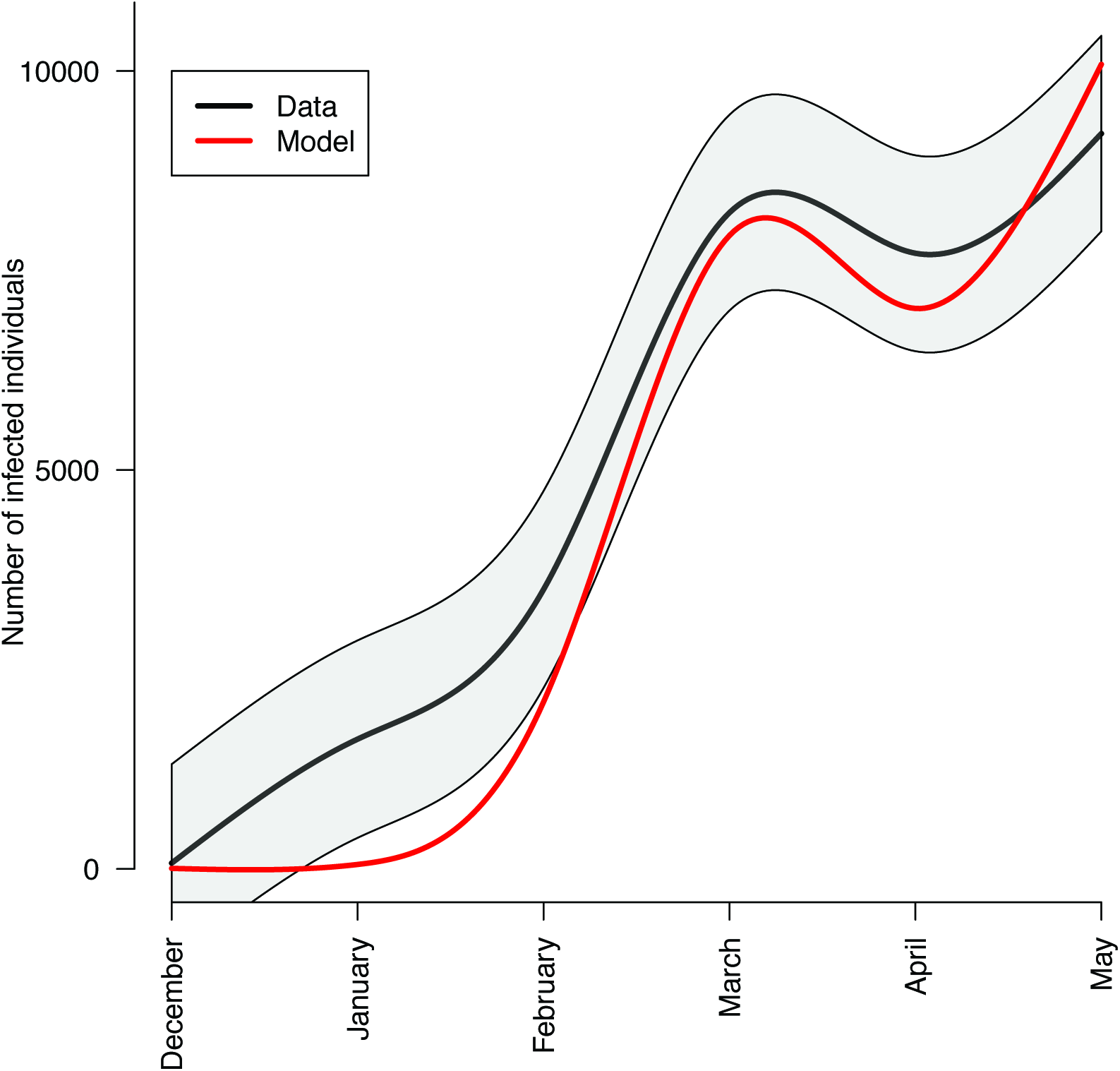
**Match between recorded epidemiological data (black) and the most parsimonious mathematical model including mosquito abundance and expressed need for protection on Twitter (red line). Estimated transmission parameters are x_0_= 3.76 10-4, x_1_=0.295, x_2_=0.644. See table 1 for model selection details.**

## Discussion

In this paper, we aimed to identify the main drivers of the Chikungunya outbreak that occurred in Martinique in 2014. We have demonstrated that the temporal dynamics of virus transmission is jointly determined by mosquito abundance as well as by human behavior as inferred from the feelings expressed on the online social network Twitter. We have also shown that the spatio-temporal dynamics of the outbreak follow a classic travelling wave pattern through space in two different directions (northwards and southwards).

While mosquito abundance is an expected explanation for the temporal dynamics of the outbreak, the importance of human behavior on the initial stages of an outbreak are less often taken into account, let alone quantified. To the best of our knowledge, this is first time that human behavior has been quantified through social media for a vector-borne disease outbreak. It is consistent with the increasing amount of evidence regarding the role of human behavior in pathogen transmission (14, 34). Moreover, our analysis shows that the influence of human behavior is not constant through time, suggesting that communication campaigns have different impacts according to the timing of the operation. This calls for a coordinated outbreak response from public health authorities, municipalities and other partners involved in source reduction, communication campaigns and use of chemical insecticide.

The spatio-temporal spread of the outbreak was shown to follow a two-waves pattern originating from the capital (Fort-De-France), which is also the place where the outbreak started. Nevertheless, the north and south waves proceeded differently. In the north, where the road network seems weakly connected, only the epidemic peak is associated with distance from Fort-De-France, suggesting that human movement acts as a seed for local epidemics. Conversely, the other traveling wave in the south, where the road network looks much more connected, is associated with similarity between the whole time series instead of the epidemics peak. We can therefore suggest that the dense road network could allow more exchange of infectious individuals that would increase the inter-dependence of pathogen dynamics between localities. While this hypothesis is difficult to demonstrate, we conducted a complementary theoretical approach (supplementary materials S4) showing that this explanation could be plausible. The suggestions that the topology of connections between human populations affects the spatial diffusion dynamics of diseases epidemics is in line with results from theoretical studies on the role of network structures on infectious disease spread (35).

As for any study focusing on epidemiological data, we have made several assumptions that deserve discussion. First, our epidemiological data relies on a sentinel network, and this non-exhaustivity could bring some uncertainty to the validity of our assumptions. Nonetheless, almost 20% of the medical doctors working on the island serve as sentinel doctors, as compared with the 1% of sentinel doctors in metropolitan France (36), providing support for the relevance and quality of the data we used in our model. Second, we have assumed that these epidemiological data contained the localities where individuals had been infected, although it instead contained the localities where the offices of sentinel doctors are located. Nevertheless, this bias is generally assumed for the study of spatio-temporal dynamics of pathogens and is not expected to play a large role on such analysis(1, 9). Finally, the low specificity of the Chikungunya symptoms, combined with an unknown number of infected individuals who did not enter the surveillance system for different reasons, can potentially lead to an underestimation of the real incidence. Nevertheless, there is no reason to believe that this bias fluctuated through time. Therefore, this might have a possible quantitative impact on our results but it should not qualitatively change our conclusions.

The use of feelings expressed on social networks, even if extremely interesting, can potentially introduce a bias since social networks are used only by a small – but rapidly growing - subset of the population (37). Recent studies have shown that the demographic makeup of social media users become increasingly less biased (38). In Martinique during the sampling period, the source of the tweets used in this study is likely to have come from a non-representative population sample. There are however no good quantitative data to suggest that epidemic awareness and the felt need for protection are very different in different population groups. Previously, the use of such digital epidemiology methods (39, 40) has been shown to reflect quite correctly the epidemiological dynamics of such emerging pathogens (39), suggesting that this surrogate can be integrated into mechanistic models involving human host behavior (14). A careful comparison of this approach against large-scale surveys, if representing the gold-standard to infer human sentiments, is lacking right now.

Human behavior is increasingly recognized as an important factor for pathogen spreading. The recent example of Ebola outbreaks in Western Africa has seen many dramatic examples of this, underlining how epidemiological response strategies need to consider this component in order to be maximally efficient (41). While the role of human behavior has long been suggested as a driver for emerging infections (42), the emergence of online social networks, now widely used throughout the world, opens new opportunities to assess it quantitatively (43). Our finding that the most parsimonious model for temporal dynamics includes Twitter activity as an anticipated indicator of its impact highlights that analysis of Twitter messages could potentially offer to public health authorities a tool to measure or even predict fluctuations in protective behavior seen in the population. Such quantification, when combined with individuals’ movements and other biotic and abiotic factors known to influence pathogen transmission, can contribute to optimize the efficiency of public health strategies (44) designed to mitigate the spread of emergent pathogens.

## Acknowledgments

We thank the “Agence Nationale de Recherche sur le Sida et les hépatites virales” (ANRS) and the “Institut de Microbiologie et Maladies Infectieuses” (IMMI) for financial support.

